# Bridging the lipid divide: archaeal ESCRT-III binds phosphoinositol and polarises the cytokinetic membrane

**DOI:** 10.64898/2026.04.15.718656

**Authors:** Alice Cezanne, Tina Drobnič, Kerstin Fiege, Yin-Wei Kuo, Joe Parham, Nicole J. Bale, Sherman Foo, Jan Löwe, Laura Villanueva, Buzz Baum

**Author notes:** These authors contributed equally to this work (Tina Drobnič, Kerstin Fiege). Corresponding authors **Email:** &. **Author Contributions:** A.C. and B.B. designed the research; A.C., T.D., K.F., Y-W.K., J.P. and S.F. performed research; A.C., K.F., J.P. and N.J.B. analysed data; A.C. wrote the paper; all authors revised the paper.

## Abstract

All cells remodel their membranes to divide. The highly conserved ESCRT-III system forms contractile polymers which, through direct interactions with membrane lipids, remodel membranes across the tree of life. In exploring how ESCRT-III divides the chemically and structurally unique archaeal membrane, we reveal that the homologue CdvB1 is required for the establishment of a distinct membrane domain within the division bridge of *Sulfolobus acidocaldarius*, associated with an accumulation of membrane-spanning inositol phosphate lipids. We show that CdvB1 associates with phosphoinositides *in vitro* and that this interaction aids cytokinesis *in vivo*. Together, we suggest that although eukaryotes inherited their membrane lipids from bacteria during eukaryogenesis, key features of the ESCRT-III:membrane interface that allow these polymers to bind, organise, and remodel eukaryotic membranes, may originate in archaea.

## Introduction

Across the tree of life, cells remodel their membranes during division. In cells that lack a cell wall, cytokinesis relies on contractile polymer networks that interact with the internal face of the membrane ^1,2^. As the division furrow ingresses, a continuous protein-membrane interface must be maintained to ensure efficient membrane constriction and abscission. In eukaryotic cells, the final stages of this process are carried out by the AAA-ATPase Vps4 ^3,4^ and ESCRT-III polymers that interact with negatively charged lipids ^5^, including specific phosphoinositides ^6–8^, insert amphipathic N-terminal helices into the membrane ^9–11^ and, in this way, perturb local lipid organisation and order ^7,12,13^.

Sequence and structural analyses have revealed that ESCRT-III predates the last universal common ancestor ^14^. In some modern archaea, cytokinesis is entirely driven by homologues of ESCRT-III and Vps4 ^15,16^. One of the best studied examples of this is *Sulfolobus acidocaldarius*, an archaeon with optimum growth around pH 3 and 75°C, which divides by the successive assembly and Vps4-mediated disassembly of the ESCRT-III polymers CdvB, CdvB1 and CdvB2 ^17–21^.

While the ESCRT-III machinery is conserved between archaea and eukaryotes, the membrane it acts on is not. Eukaryogenesis is widely accepted to have involved the merger of an archaeal cell related to modern day Asgard archaea (also ‘Promethearchaeota’), and a bacterial cell related to modern day Alphaproteobacteria. During this process, eukaryotes inherited many membrane proteins from their archaeal ancestor, including ESCRT-III, the translocon and V-ATPase ^22,23^. However, the membrane lipids of all living eukaryotes are synthesised by enzymes inherited from bacteria ^24,25^.

This is significant, as the lipids of archaea and bacteria are profoundly different, a phenomenon termed the “lipid divide”. Bacterial and eukaryotic lipids, formed by sn-glycerol-3-phosphate backbones linked via ester linkage to fatty acid chains, organise into bilayers. Archaeal lipids, however, formed by sn-glycerol-1-phosphate backbones linked via ether linkages to saturated, methyl-branched isoprenoid chains, can exist in either bilayer or monolayer form ^24,26,27^. The monolayer form comprises bi-polar membrane spanning lipids, synthesised from two bilayer lipids, and makes up the majority of the membrane in *S. acidocaldarius* and other thermoacidophiles ^28–30^. Lateral organisation is regulated by the addition of cyclopentane rings to the monolayer lipid core ^31,32^ but apolar membrane components, such as squalene or respiratory quinones, have also been speculated to play a role ^33,34^. Headgroups are shared across all domains of life, and many extremophile archaea, such as *S. acidocaldarius*, use polyglycosylated lipids to protect cell integrity ^24,35^. By virtue of these chemically and structurally unusual lipids, archaeal membranes are highly impermeable ^36^, but relatively non-deformable ^37^, allowing these cells to maintain a fluid membrane and survive in extreme environmental conditions.

To understand how a conserved membrane remodeling machinery, such as ESCRT-III, can act on either side of the lipid divide, we investigated the nature of the interface between the archaeal membrane and the contractile ESCRT-III polymers that drive cytokinesis in *S. acidocaldarius*, CdvB1 and CdvB2. In doing so, we show that, despite the profound changes in lipid chemistry that accompanied eukaryogenesis, key interactions between ESCRT-III and the lipid membrane have remained relatively unchanged.

## Results

### CdvB1 establishes the formation of a cytokinetic domain with locally altered membrane content

For this investigation we used a dominant negative Vps4 mutation, Vps4^E209Q^, which arrests cells during constriction, without blocking cell growth or progression through the cell cycle. This leads to accumulation of large cells with an exaggerated cytokinetic bridge that can be easily studied by light microscopy ^20^. We imaged the membranes of live Vps4^E209Q^ cells under normal growth conditions (75°C, pH 3) and observed distinct partitioning of lipid dyes (Fig 1A, S1A). CellMask Plasma Membrane Stain (Invitrogen C10046; hereafter CellMask) was excluded from a broad region of the cytokinetic bridge, while NileRed was enriched. These staining patterns were not due to interactions between the dyes, as the same result was seen with CellMask alone (Fig S1A). In addition, we observed a similar, pattern of CellMask staining in a subset of actively constricting MW001 control cells, albeit less striking than in Vps4^E209Q^ (Fig 1B, S1B, Movies S1-3). In MW001 cells, a gap in CellMask staining appeared at the midzone ~6 minutes before the start of constriction, implying that it is associated with division ring assembly ^38^ rather than membrane curvature, and was not a peculiarity of Vps4^E209Q^ expression.

**Figure 1.**
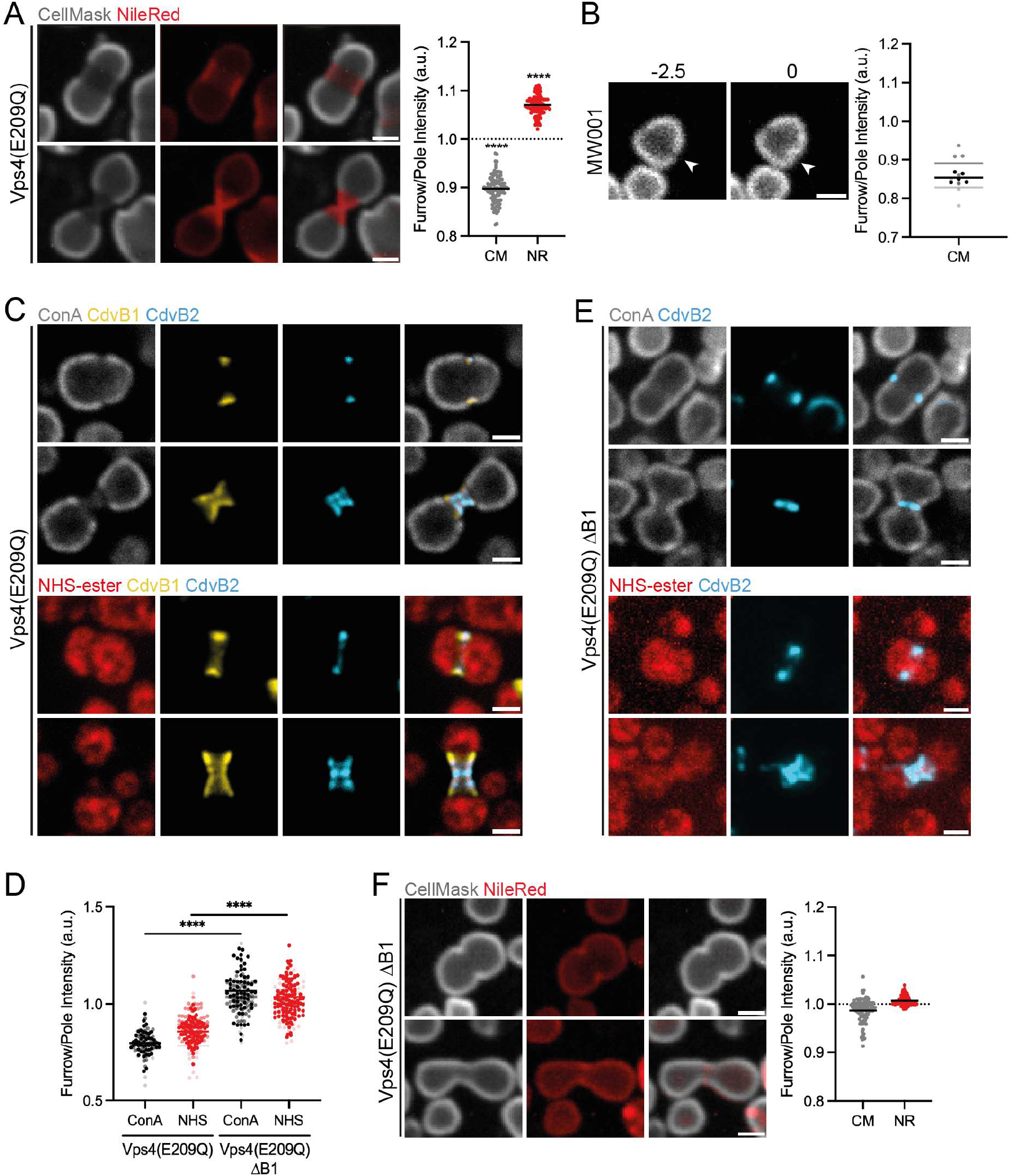
CdvB1 forms a specialised membrane domain at the cytokinetic furrow. (**A**) Live Vps4^E209Q^ cells stained with CellMask Deep Red Plasma Membrane Stain and Nile Red at 75°C after 8 hours of arabinose induction (N = 5) (*left*) and the ratio of Furrow to Pole intensity for each stain (solid line denotes mean) (*right*). Significance values were derived using a one sample ratio *t* test to test whether the values were significantly different from 1 (n = 100). (**B**) Live cell imaging of an MW001 cell stained with CellMask Deep Red Plasma Membrane stain at 75°C at 2.5 minutes before the onset of constriction (−2.5; future position of furrow marked with an arrow) and at the onset of constriction (0; initial furrow ingression marked with an arrow) (*left*) and the ratio of Furrow to Pole intensity (line denotes mean, N = 3, n = 5) (*right*). (**C**) Representative immunofluorescence images of ethanol fixed Vps4^E209Q^ after 8 hours of arabinose induction stained for ConA, CdvB1 & CdvB2 (*top*) or NHS-ester, CdvB1 & CdvB2 (*bottom*). (**D**) Ratio of Furrow to Pole intensity for ConA & NHS-ester stains (line denotes mean). Significance values were derived using Welch’s *t* test (N = 3, n > 50). (**E**) Representative immunofluorescence images of ethanol fixed Vps4^E209Q^ ΔB1 after 8 hours of arabinose induction stained for ConA & CdvB2 (*top*) or NHS-ester & CdvB2 (*bottom*). (**F**) Live Vps4^E209Q^ ΔB1 stained with CellMask Deep Red Plasma Membrane Stain and Nile Red at 75°C after 8 hours of induction (N = 3) (*left*) and the ratio of Furrow to Pole intensity for each stain (solid line denotes mean, n = 91) (*right*). Scale bar = 1µm. Each image represents a single z-slice. Merged images were generated using the maximum values across all channels to better visualise position of staining relative to each other (ImageJ Composite Max display mode).

We next compared the pattern of lipid dye staining in Vps4^E209Q^ cells with the position of the contractile ESCRT-III proteins, CdvB1 and CdvB2, by immunofluorescence. While CdvB2 localised to discrete bands at the narrowest regions of the bridge, CdvB1 almost homogenously coated a broad central region, similar in size and shape to the CellMask-depleted region (Fig 1C). This region was also poorly labelled with two fluorescently conjugated labels: ConA (labels glycosylated surface proteins and lipids), and NHS-ester (labels all accessible amines) (Fig 1C & D), suggesting that it may be depleted of transmembrane proteins. To test this, we overexpressed a single-pass transmembrane protein, the Cdc48 homologue Saci_0877 (HA-0877) and confirmed that its expression did not perturb division (Fig S2A). Immunofluorescence revealed that HA-0877 was clearly excluded from the region of the membrane occupied by ESCRT-III rings (Fig S2B).

As the patterns of membrane and protein staining closely matched the footprint of CdvB1, we repeated this investigation in a CdvB1 deletion strain expressing Vps4^E209Q^ (Vps4^E209Q^ ΔB1). Strikingly, both ConA and NHS-ester labelled the cytokinetic bridge in Vps4^E209Q^ ΔB1 cells, even though CdvB2 remained and, in some cases, occupied a broad region of the membrane (Fig 1E). In live cell imaging, the bridges of Vps4^E209Q^ ΔB1 cells failed to exclude CellMask (Fig 1F, S1C) and had a less polarised Nile Red signal. This suggests that, in the presence of CdvB1, the membrane of the cytokinetic bridge in *S. acidocaldarius* has a distinct protein and lipid content.

### Inositol phosphate monolayer lipids are produced in a CdvB1-dependent manner in constricting cells

To characterise this domain, we investigated to what extent membrane lipid composition changes in dividing *S. acidocaldarius* cells. We performed intact polar lipid analysis of Vps4^E209Q^ cells (60% of cells display cytokinetic bridges compared to <4% in MW001 control ^39^), and compared this to Vps4^E209Q^ ΔB1 cells. The analysis identified a set of respiratory quinones and different classes of polar lipids consisting of a core lipid conjugated to one or more headgroups (Table 1, Fig S3A, Tables S1-3). The core lipid was either bilayer-forming C20 dialkyl glycerol diether (DGD, hereafter ‘bilayer lipid’) or monolayer-forming C40 glycerol dialkyl glycerol tetraether (GDGT, hereafter ‘monolayer lipid’) containing between zero and six cyclopentane rings. The most abundant headgroups were inositol phosphate (IP), monohexose (MH), dihexose (DH) and dihexose & inositol phosphate (DH-IP). Each class may represent multiple headgroup configurations or topologies (e.g. symmetric dihexose lipids or *Sulfolobus* specific sugar groups, e.g. calditol ^40^). Representative structures, based on MS^2^ fragmentation patterns, and relative abundances are shown in Fig S3A & B.

**Table 1.** Relative abundances of lipid classes identified in MW001, Vps4^E209Q &^ Vps4^E209Q^ ΔB1 as a fraction of total polar lipids. Values are given as Mean ± SD (N = 3).

| Lipid core | Headgroup | MW001 (%) | Vps4<sup>E209Q</sup> (%) | Vps4<sup>E209Q</sup> ΔB1 (%) |
| --- | --- | --- | --- | --- |
| Bilayer (DGD) | none | 0.22 ± 0.02 | 0.62 ± 0.06 | 0.38 ± 0.09 |
|  | IP | 16.67 ± 1.14 | 12.56 ± 1.86 | 15.71 ± 8.02 |
| Monolayer (GDGT) | none | 21.49 ± 1.26 | 6.32 ± 0.66 | 16.39 ± 2.42 |
|  | MH | 0.77 ± 0.09 | 0.31 ± 0.02 | 1.99 ± 0.27 |
|  | DH | 22.59 ± 1.48 | 12.24 ± 1.90 | 29.44 ± 4.79 |
|  | IP | 35.78 ± 2.63 | 44.16 ± 3.25 | 32.92 ± 1.22 |
|  | DH-IP | 2.49 ± 0.82 | 23.79 ± 3.80 | 3.15 ± 0.75 |

We first examined whether the cytokinetic domain was accompanied by any changes to the hydrophobic core of the membrane. Notably, none of the samples showed alterations to the ratio of bilayer to monolayer lipids (Fig 2A). There was a marked increase in the abundance of quinones relative to total polar lipids in Vps4^E209Q^ but not Vps4^E209Q^ ΔB1 cells (Fig S3C). Most significantly, Vps4^E209Q^ cells showed dramatically reduced cyclisation of monolayer lipids, that was not observed in Vps4^E209Q^ ΔB1 cells (Fig 2B; Fig S3D). This suggests that the existence of this CdvB1-dependent cytokinetic domain is accompanied by changes in lateral lipid organisation.

**Figure 2.**
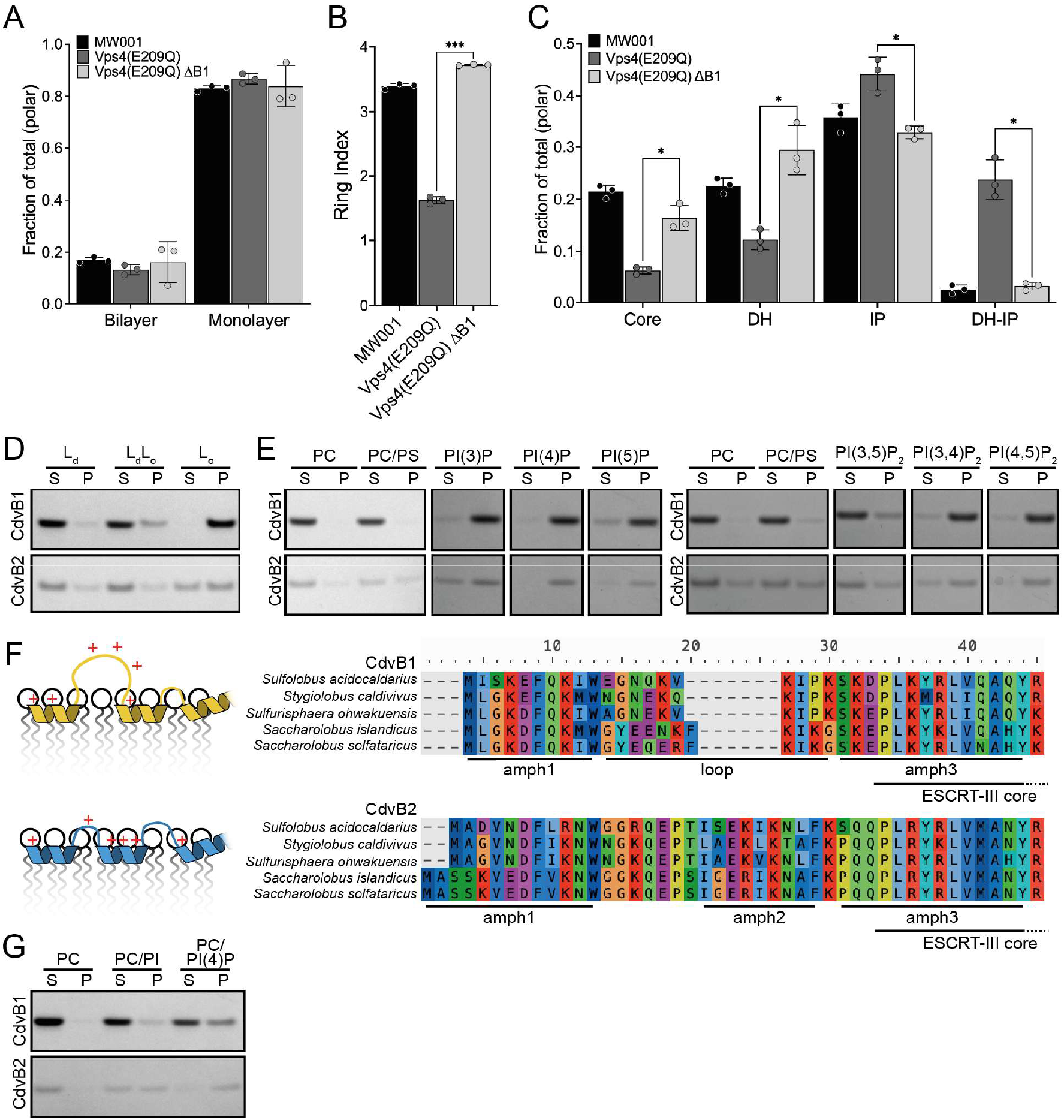
Inositol phosphate lipids are produced during constriction in a CdvB1-dependent manner and recruit CdvB1 and CdvB2 *in vitro*. (**A**) Fractions of bilayer and monolayer lipids in MW001, Vps4^E209Q^ & Vps4^E209Q^ ΔB1 as a percentage of total polar lipids. Significance values were derived using Welch’s *t* test (N = 3). Mean plotted with error bars denoting ± SD. (**B**) Ring index of monolayer lipids in MW001, Vps4^E209Q &^ Vps4^E209Q^ ΔB1. Ring index is a weighted average of ring numbers on monolayer lipids and was calculated by multiplying the relative abundance of each lipid class by the number of rings present in that class (i.e. (abundance_0 × 0) + (abundance_1 × 1) + (abundance_2 × 2) etc.). Significance values were derived using Welch’s *t* test (N = 3). Mean plotted with error bars denoting ± SD. (**C**) Monolayer lipid headgroup distribution in MW001, Vps4^E209Q^ & Vps4^E209Q^ ΔB1 as a percentage of total monolayer lipids. Significance values were derived using Welsch’s *t* test (N = 3). Mean plotted with error bars denoting ± SD. (**D**) In vitro pelleting of CdvB1 and CdvB2 in L_d_ (PC:SM:CH 80:15:5), L_d_/L_o_ (PC:SM:CH 37.5:37.5:25), and L_o_ (SM:CH 60:40) liposomes. (**E**) In vitro pelleting of CdvB1 and CdvB2 in PC/PIP (80:20 PC:PI(3)P/PI(4)P/PI(5)P) and PC/PIP2 liposomes (80:20 PC:PI(3,4)P_2_/PI(3,5)P_2_/PI(4,5)P_2_). (**F**) Schematic representation of membrane interaction site and basic amino acid distribution (*left*) and alignment of archaeal CdvB1 and CdvB2 N-termini (*right*). (**G**) In vitro pelleting of CdvB1 and CdvB2 in PC, PC/PI (95:5) and PC/PI(4)P (95:5) liposomes. For all: S = supernatant & P = pellet. NB: in the pelleting assay, a pre-spin to remove protein aggregates resulted in lower CdvB2 signal.

Since ESCRT-III proteins bind negatively charged lipids ^5,11,41–43^, we were especially interested in potential sources of negative charge that could be involved in the association of CdvB1 with the membrane. The analysis did not identify classic negatively-charged headgroups such as serine or phosphoglycerol, as *S. acidocaldarius* lacks the genes to produce them ^44^. Of the classes identified, only inositol phosphate containing lipids (IP and DH-IP) unequivocally display a negative charge: the phosphate group between the inositol and the glycerol backbone. Intriguingly, unlike in eukaryotes, archaeal biosynthesis of these lipids involves the addition of a 1L-myo-inositol 1-phosphate to cytidine diphosphate-DGD (CDP-DGD), generating archaetidylinositol(3)phosphate (AI(3)P), a bilayer lipid analogous to the eukaryotic phosphatidylinositol(3)phosphate (PI(3)P) (Fig S3E) ^45^. While archaeal cell extracts demonstrate phosphatase activity generating archaetidylinositol (AI) from AI(3)P, no archaetidylinositol phosphatases or kinases have been identified to date ^46^.

In our analysis, inositol phosphate lipids made up the majority of the membrane in *S. acidocaldarius* (See Table 1; Fig S3B). Vps4^E209Q^ cells showed a dramatic increase in the relative abundance of DH-IP monolayer lipids, an otherwise minor component of the membrane, and a modest increase in IP monolayer lipids (Fig 2C). We also observed a reduction in the relative abundance of core and DH monolayer lipids. Strikingly, all of these changes depended on the presence of CdvB1, suggesting the possibility of direct interactions between ESCRT-III and inositol phosphate lipids during archaeal cytokinesis.

### CdvB1 binds tightly packed lipid membranes and phosphoinositide lipids in vitro

To explore this potential interaction, we turned to *in vitro* reconstitution. Recombinant CdvB1 and CdvB2 were purified and incubated with liposomes composed of eukaryotic lipids, as synthetic archaeal-type lipids are not yet available. The strength of protein-membrane interaction was evaluated by a pelleting assay, comparing the amount of protein in the lipid bound (pellet) and unbound (supernatant) fractions after centrifugation.

Since these are bacterial-type lipids, we first tested the contribution of the hydrophobic core. We assessed binding to liposomes that lacked negatively charged lipids and consisted of only the liquid disordered phase (L_d_; loosely packed), liquid ordered phase (L_o_; tightly packed), or a mixture of both phases (L_d_/L_o_) (Fig 2D). CdvB1 showed extremely strong binding to L_o_, weak binding to L_d_/L_o_ and no binding to L_d_ membranes. CdvB2, included as a control, was able to bind all three, but showed weaker binding to L_o_ membranes compared to CdvB1. This suggests that CdvB1 interacts less strongly with the hydrophobic core of the membrane than CdvB2, but can stably associate with liposomes in the absence of negative charge, provided lipids are tightly packed, i.e. conditions more akin those found in archaeal monolayer membranes.

Next, we investigated the contribution of lipid headgroups and negative charge. To address whether electrostatic interactions can occur with inositol phosphate lipids, we compared the recruitment of CdvB1 and CdvB2 to disordered phosphatidylcholine (PC) membranes containing either phosphatidylserine (PS) or phosphatidylinositolphosphate (PIP) as a source of negative charge. CdvB1 showed no binding to PC and very little binding to PC/PS membranes, whereas CdvB2 was recruited to both (Fig 2E). Replacing PS with PIP however, allowed CdvB1 to bind strongly and with unexpected specificity. CdvB1 bound most strongly to PI(3)P and PI(4)P membranes and also bound strongly to PI(3,4)P_2_ and PI(4,5)P_2_. CdvB2 displayed a broadly similar preference, with the exception of weaker binding to PI(3)P and PI(3,4)P_2_.

To better understand how CdvB1 and CdvB2 interface with the membrane, we examined their membrane binding domains (Fig 2F). The N-terminal regions of CdvB1 and CdvB2 contain the entire membrane interaction interface, consisting of two properties: the insertion of amphipathic helices into the hydrophobic core and electrostatic interactions with lipid headgroups ^7,11,42,47^. While CdvB2 has three N-terminal amphipathic helices, the CdvB1 N-terminus contains only two predicted amphipathic helices, connected by a positively charged ~10 amino acid loop ^11^. This could allow CdvB1, but not CdvB2, to interact with charged groups that sit above the plane of the membrane. To test this, we investigated recruitment to membranes containing phosphatidylinositol (PI) which lacks the raised phosphate group. While CdvB2 bound very strongly to membranes containing PI, CdvB1 did not (Fig 2G). Together, this suggests that archaeal ESCRT-III proteins can interact with phosphoinositides, and that CdvB1 is highly sensitive to the height of the charged group.

### CdvB1 and inositol phosphate lipid interactions facilitate membrane constriction in vivo

Since the *S. acidocaldarius* membrane contains no other obvious sources of negative charge, *in vivo*, interactions between inositol phosphate lipids and ESCRT-III may play a role in cytokinesis. In *Sulfolobus*, inositol phosphate lipids are generated by the archaetidylinositolphosphate synthase (AIPS) (Fig S3E). AIPS is a ~21kDa transmembrane protein, structurally homologous to both the bacterial phosphatidylinositolphosphate synthase and the eukaryotic phosphatidylinositol synthase ^45^. AIPS is essential in *Saccharolobus islandicus* ^48^ and we could not knock it out in *S. acidocaldarius*. Thus, we tested the impact of AIPS overexpression on cell division in MW001 control (AIPS-HA) and CdvB1 deletion (AIPS-HA ΔB1) backgrounds.

Immunofluorescence revealed that AIPS-HA is homogenously distributed in the membrane but, like other transmembrane proteins, excluded from the cytokinetic furrow during constriction. (Fig 3A). Cells expressing AIPS-HA did not show any obvious division phenotypes and exhibited normal levels and profiles of CdvB1 and CdvB2 as assessed by flow cytometry (Fig S4) and immunofluorescence (Fig 3B). Live cell imaging revealed that the rate of furrow constriction was significantly increased upon expression of AIPS-HA (Fig 3C). Strikingly, this effect was abolished in ΔB1 cells, which displayed a reduced rate of constriction that was indistinguishable between control and AIPS-HA expressing ΔB1 cells (Fig 3D).

**Figure 3.**
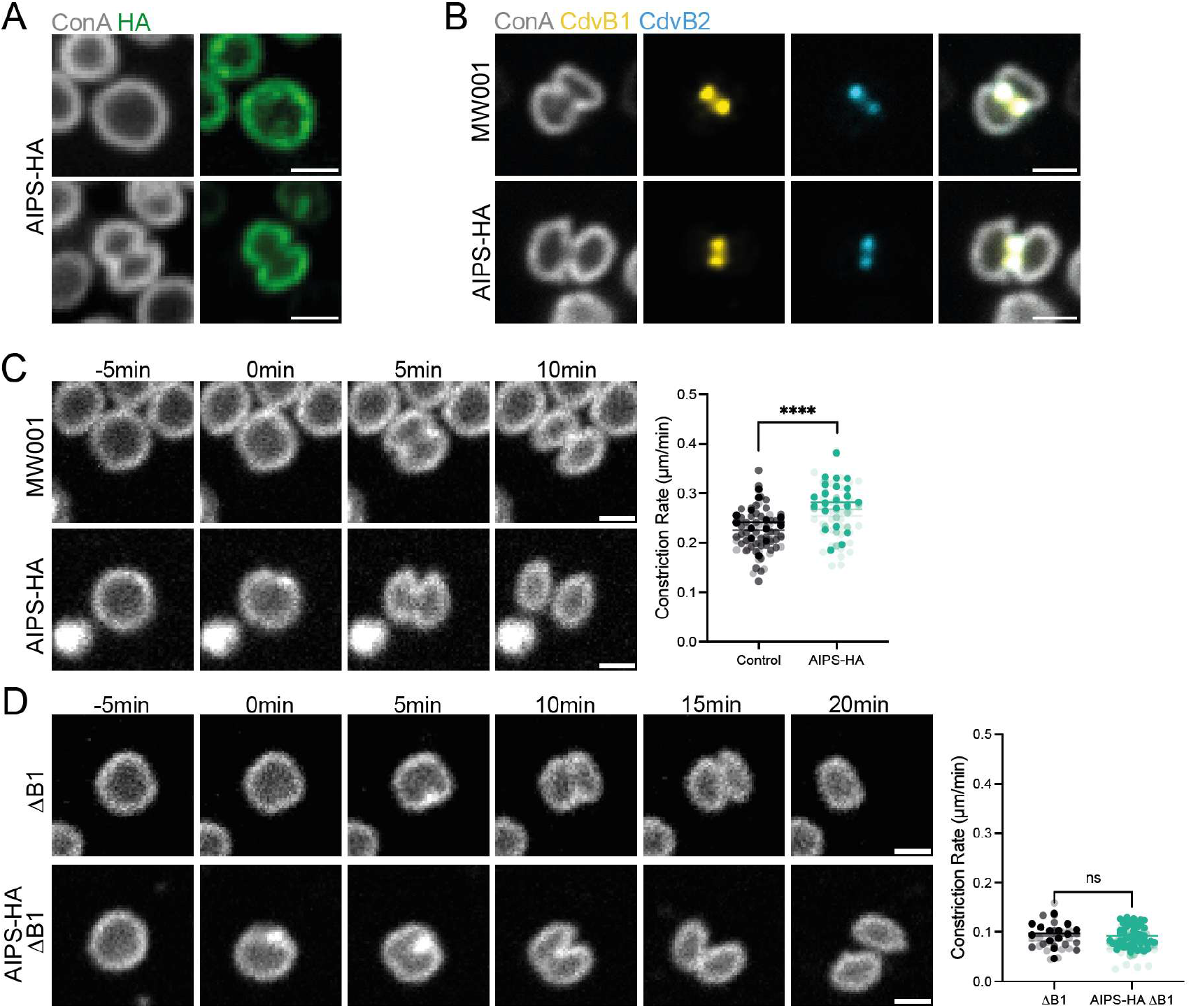
Overexpression of the archaetidylinositolphosphate synthase enhances ESCRT-mediated membrane constriction in a CdvB1 dependent manner. (**A**) Immunofluorescence images of formaldehyde fixed AIPS-HA cells after 4 hours of arabinose induction, stained for ConA & HA. (**B**) Immunofluorescence images of ethanol fixed MW001 and AIPS-HA cells after 4 hours of arabinose induction stained for ConA, CdvB1 & CdvB2. (**C**) Time lapse images of cell division in MW001 empty vector control (MW001 EV) and AIPS-HA cells after 4 hours of arabinose induction, stained with CellMask Deep Red Plasma Membrane stain at 75°C (*left*) and calculated constriction rate for each strain (*right*). Significance values were derived using Welch’s *t* test (N = 3, n > 15). (**D**) Time lapse images of cell division in ΔCdvB1 empty vector control (ΔB1 EV) & AIPS-HA ΔB1 cells after 4 hours of arabinose induction, stained with CellMask Deep Red Plasma Membrane stain at 75°C (*left*) and calculated constriction rate for each strain (*right*; line denotes mean). Significance values were derived using Welch’s *t* test (N = 3, n > 15). Scale bar = 1µm. Merged images were generated using the sum of values across all channels (ImageJ Composite display mode).

## Discussion

In this study, we make the surprising finding that the membrane at the cytokinetic bridge in *S. acidocaldarius* cells is distinct from the rest of the cell. Discrete, functionalised membrane domains have been shown to contribute to vital cellular functions in bacteria and eukaryotes ^49,50^, but, while their existence in archaea has long been a source of speculation, have not yet been observed *in vivo* or *in vitro* ^37,51,52^. Here, we show that the formation of a cytokinetic domain depended nearly entirely on the presence of the ESCRT-III homologue CdvB1, which forms an extended polymer across the division bridge. We propose here that CdvB1 organises membrane contents to facilitate robust and efficient constriction.

Our data suggest that this activity arises from the interface of CdvB1 with membrane lipids. The CdvB1 N-terminus displays two putative amphipathic helices, connected by a flexible charged loop. We propose that the amphipathic helices insert into the membrane while the flexible loop is available for electrostatic interactions above the plane of the membrane. Compared to CdvB2, this architecture allows the helices of CdvB1 more freedom of movement and reduces their footprint in the hydrophobic core, minimising perturbation of the membrane. We speculate that this allows CdvB1 to occupy a broad region of the membrane, over a range of curvatures, and that large, continuous assemblies of CdvB1 polymer could sterically result in the observed exclusion of transmembrane proteins from the cytokinetic domain. While we show that transmembrane proteins are broadly excluded from the cytokinetic domain, we previously showed that the proteinaceous S-layer lattice, formed in part by the glycosylated, single-pass transmembrane protein SlaB, is intact across the division bridge and necessary for efficient cytokinesis ^53^. We suggest that, due to a short cytoplasmic tail and external anchoring, SlaB may be retained in the cytokinetic domain while other transmembrane proteins are not. Excluding transmembrane proteins may support a continuous ESCRT-membrane interface and more efficient inward curvature generation. Further, the interface between the ESCRT-rich cytokinetic bridge and the flanking transmembrane protein-rich membrane may aid remodeling and scission by forming a boundary differential tension, as seen in other systems ^54–56^.

Analysis of the lipid composition in *S. acidocaldarius* showed that the CdvB1-dependent cytokinetic domain is accompanied by an increase in the relative abundance of inositol phosphate monolayer lipids. *In vitro*, CdvB1 interacts weakly with sources of negative charge at or below the plane of the membrane but strongly with phosphoinositides, where charged groups are raised above the plane of the membrane. The lipid analysis also revealed the cytokinetic domain is accompanied by a reduction in monolayer lipid cyclisation, which increases membrane fluidity and polar headgroup height ^57–59^. In this way, the cell might balance the need for an impermeable yet more flexible membrane during cytokinesis in a hot, acidic environment. Decreasing cyclisation raises the headgroup from the plane of the membrane and could enhance the presentation of inositol phosphate headgroups for CdvB1 interaction. Finally, *in vivo*, overexpression of the enzyme that produces inositol phosphate lipids enhances the rate of furrow constriction, but only in the presence of CdvB1, supporting a role for association of these lipids with CdvB1 during cytokinesis.

This study suggests unexpected commonalities between archaeal and eukaryotic membranes. In eukaryotes, the membrane interface of ESCRT-III relies on direct interactions with negatively charged phosphoserine lipids, and sometimes specific phosphoinositides, as well as modification of lipid tail length and saturation to alter membrane fluidity and presentation of lipid headgroups ^5–10,12,13^. Here, we show that orthogonal processes and chemistries still result in similar features of the ESCRT-III:membrane interface in archaea. This suggests that during eukaryogenesis, a transition from archaeal- to bacterial-type lipid membranes may not have posed as much of a problem for ESCRT-III as previously assumed. Intriguingly, the eukaryotic biosynthesis of phosphatidylinositol does not originate in archaea, implying that the functions of these lipids, but not their biosynthetic pathways, were retained along with ESCRTs during eukaryogenesis ^60^. Our data suggests the possibility that inositol phosphate lipids had a role in protein interaction and establishment of local membrane identity, prior to the emergence of the eukaryotic endomembrane system. Future investigations, in *Sulfolobus* and species more closely related to the archaeal ancestor of eukaryotes, will reveal in how far the evolution of the endomembrane system was shaped by these pre-existing activities, or other membrane features that were, until recently, thought to be eukaryote specific.

By identifying common aspects of lipid biology, this work helps to understand how, despite radically different membrane chemistries, archaeal-derived membrane proteins are able to carry out functions in eukaryotes similar to those their progenitors carried out in archaea.

## Materials and Methods

### Cloning and transformation

To generate AIPS-HA overexpression plasmid, pSVAaraFX-HA plasmid vector was double digested with restriction enzymes NcoI and XhoI followed by gel extraction. The endogenous sequence of AIPS (saci_1541) was PCR amplified from the DSM639 genome to introduce a Gly-Gly-Gly-Ser linker flanking the C-terminal end of the gene using the forward primer: 5’-AATAATTGATAAGCGTCTTACTTATCATACCATGCTTACAAGAATACGAAAACAGTC and the reverse primer: 5’-TACGCGTAGTCCGGAACGTCATACGGGTACTCGAGTGATCCTCCACCGCTAAGGTTAAAGTAAATGTAAATGAAC (regions annealed to the saci_1541 gene are underlined). The PCR product was then inserted into the digested vector by Gibson assembly. For cloning of saci_0877^E162A/E433A^-HA expression plasmid, custom synthesized double stranded DNA fragment (gBlock, Integrated DNA Technologies) corresponding to the codon optimized saci_0877^E162A/E433A^ gene flanking with two overlapping sequences with the NcoI/XhoI digested pSVAaraFX-HA vector was used as insert and assembled by Gibson assembly. All plasmids were verified by Sanger sequencing. Plasmids were transformed into electrocompetent *S. acidocaldarius* MW001 and ΔCdvB1 using electroporation (2000 V, 25 μF, 600 ohms, 1 mm) and selected on Gelrite-Brock plates.

### Cell Culturing & Strains

*S. acidocaldarius* strains MW001, MW001 EV (MW001 with the empty vector plasmid), Vps4^E209Q^ (MW001 with Vps4^E209Q^ -6xHis), 0877 (MW001 with DN0877-HA), AIPS-HA (MW001 with AIPS-HA) and the mutant strains ΔB1 EV (ΔB1 strain with the empty vector plasmid), AIPS-HA ΔB1 (ΔB1 with AIPS-HA) Vps4^E209Q^ ΔB1 (ΔB1 with Vps4^E209Q^ -6xHis) were grown in Brock medium (pH 2.9) supplemented with 0.1% N-Z-amine and 0.2% sucrose in a shaking incubator set to 75°C. Overexpression of EV, Vps4^E209Q^ -6xHis, 0877-HA, AIPS-HA strains was induced by addition of arabinose for 4 or 8 hours at a final concentration of 0.2% (w/v). All cultures used for imaging or lipidomics were collected during exponential growth phase (OD_600nm_ of 0.15 to ~0.4). Arabinose induction was performed at an OD_600nm_ ~0.15.

### Live cell-imaging at 75°C

Live-cell imaging was performed at 75°C using the Sulfoscope chamber described previously ^61^. In short, Attofluor chambers (Invitrogen A7816) were assembled with 25-mm coverslips, washed with EtOH and H_2_O. ~300µl Brock medium was added to the chamber and left to dry at 75°C (~45-60 minutes). Chambers were washed thoroughly with fresh Brock medium, and equilibrated to 75°C in the Sulfoscope. CellMask Deep Red Plasma Membrane Stain (Invitrogen C10046), and NileRed (Invitrogen N1142) were added to 5ml of *S. acidocaldarius* cell culture at an OD_600nm_ of 0.15 to 0.3 at 75°C immediately before imaging, to a final concentration of 1µg/ml and 2.5µg/ml respectively. 400 μl of labelled cell suspension was added into the chamber for imaging. Cells were immobilized using heated, semi-solid gelrite pads (0.6% Gelrite, 0.5× BNS pH 5, and a final concentration of 20 mM CaCl_2_). Images were acquired on a Nikon Eclipse Ti2 inverted microscope equipped with a Yokogawa SoRa scanner unit and Prime 95B sCMOS camera (Photometrics). Imaging was performed with a 60× oil immersion objective (Plan Apo 60×/1.45, Nikon) heated to 60°C by an objective collar controlled by an OKOlab UNO-T stage top incubator. Immersion oil was custom formulated for high temperature imaging (maximum refractive index matching at 70°C, *n* = 1.515 ± 0.0005; Cargille Laboratories). Using the ×2.8 magnification of the SoRa unit, images were acquired at a total magnification of ×168 with 15-ms exposure time and 10% laser power. For timelapse imaging, images were acquired at intervals of 15 seconds for 2.5 hours. For CellMask and Nile Red staining visualisations, 100 frames were acquired without interval with 15-ms exposure time and 10% laser power. After acquisition, XY drift was corrected using the ImageJ plugin StackReg ^62^. CellMask and NileRed visualization in Figure 1A and 1F was achieved by taking the sum intensity of 100 frames, performed in ImageJ.

### Cell fixation and staining

Cells stained with CdvB1 & CdvB2 antibodies were fixed by stepwise addition of ice-cold absolute ethanol to a final a concentration of 70%. AIPS-HA cells stained with HA antibody were fixed with 4% formaldehyde and treated with 0.01% SDS for 20 min. Fixed cells were washed and rehydrated PBSTA (phosphate-buffered saline supplemented with 0.1% Tween 20 and 3% bovine serum albumin) and incubated with lab generated primary antibodies to CdvB, CdvB1 and CdvB2 or commercial anti-HA (ThermoFisher, 26183) overnight at 25° C and 400rpm shaking. Secondary antibody staining was performed with Alexafluor conjugated antibodies (Alexa Fluor 405 anti-Rabbit (ThermoFisher A11034); Alexa Fluor 488 anti-Chicken (ThermoFisher A11039); Alexa Fluor 546 anti-Guinea Pig (ThermoFisher A11074); Alexa Fluor 546 anti-Mouse (ThermoFisher A11030)) for 2-3 hours at 25° C and 400rpm shaking. At the same time, surface glycosylation was stained with 200 μg/mL Concanavalin A conjugated to Alexafluor 647 (ThermoFisher, C21421) and total protein distribution with NHS-ester conjugated to Alexafluor 647 (Invitrogen A20006).

### Flow cytometry

For flow cytometry, DNA was labelled with 2μM Hoechst. Cells were gated by DNA staining (UV excitation). Laser excitation wavelengths of 355, 488, 561 and 633nm were used in conjunction with the emission filters 450/50, 530/30, 586/15 and 670/14 respectively. Flow cytometry analysis was performed on BD Biosciences LSRFortessa. Side scatter and forward scatter was recorded. Analysis was performed using FlowJo v10.8.1. Levels of ESCRT-III in G2 cells were measured by taking the high population in cells with two copies of DNA as visualised by Hoechst staining.

### Fixed cell imaging

Lab-Tek chambered slides (Thermo Fisher Scientific, 177437PK) were coated with 2% polyethyleneimine (PEI), 30min at 37° C. Coated chambers were washed with Milli-Q water before 200µl cell suspension was added and spun for 1 hour at 750 relative centrifugal force (RCF). Cells were imaged in Lab-Tek chambered coverslip using a Nikon Eclipse Ti2 inverted microscope equipped with a Yokogawa SoRa scanner unit and Prime 95B sCMOS camera (Photometrics). Images were acquired with a 100× oil immersion objective (Apo TIRF 100×/1.49, Nikon) using immersion oil (immersion oil type F2, Nikon). A total magnification of 280× was achieved using the 2.8× magnification lens in the SoRa unit. Images were acquired with 200-ms exposure time for labelled proteins and 500-ms exposure time for DNA stains, with laser power set to 20%. *z*-axis data were acquired using 6 or 10 captures with a 0.18 to 0.22-μm step (6 captures for NHS-ester staining, 10 captures for everything else). Prior to image analysis, chromatic shift was corrected in ImageJ based on a calibration image of Tetraspeck beads (Invitrogen T14792) acquired using the same settings as above.

### Image analysis

Quantification of signal intensities at the furrow vs pole was performed in ImageJ by dividing the average value of 95^th^ percentile pixel values from an equally sized ROI at the pole and furrow. For live cell images the furrow ROI was defined using NileRed staining and for immunofluorescence images the furrow ROI was defined using CdvB2 staining for Vps4^E209Q^ strains and CdvB1 signal for 0877-HA & ATPA-HA. Constriction rate analysis was performed on live cell timelapses by manually defining the first and last frames of constriction to calculated the time taken during constriction and diving this by the cell diameter (measured along the cytokinetic furrow prior to the onset of constriction). As indicated in the figure legends, merged images were generated in ImageJ using either the Composite Max mode which displays the maximum values across all channels (Fig. 1 & S1) or the traditional Composite mode which displays the sum of all values across channels (Fig. 4). We found the Composite Max mode to better show that CdvB1 staining occupies the full area from which ConA and NHS-ester staining is excluded, while CdvB2 occupies a discrete and more central area of the cytokinetic furrow. To visualise colocalization, as in Figure 4, the traditional sum Composite display was used.

### Lipid analysis

Lipid analysis was performed on 4 and 8hr inductions of MW001 and Vps4^E209Q^ strains, however no further changes in lipid profiles were observed upon longer induction. For lipid analysis, cells were collected on muffled glass fibre filters (GF-75 47 mm diameter 0.3 µm pore size, Advantec), frozen at −80°C, and freeze-dried prior extraction. Intact polar lipids (IPLs) were extracted using a modified Bligh-Dyer protocol ^63,64^. In brief, samples were ultrasonicated twice in methanol (MeOH)/dichloromethane(DCM)/phosphate buffer (2:1:0.8, v/v/v). The supernatants were collected and phase separated by adding additional DCM and phosphate buffer to a final ratio of 1:1:0.9 (v/v/v). The organic phase was collected and the aqueous phase re-extracted twice with DCM. These steps were repeated on the residual material but with MeOH/DCM/trichloroacetic acid (2:1:0.8, v/v/v). The combined DCM layers were dried under a stream of N_2_ gas and stored at −20°C. The dried extracts were reconstituted in a MeOH/DCM mix (9:1, v/v) containing an internal standard, filtered through a 0.45μm RC filter (4 mm diameter, Grace Alltech) and subsequently analysed on an Agilent 1290 Infinity I UHPLC System coupled to a Q-Exactive Orbitrap mass spectrometer (Thermo Fisher Scientific). An Acquity BEH C18 column (Waters, 2.1 × 150 mm, 1.7 μm), maintained at 30°C, using a flow rate of 0.2 mL min−1 was used for separation. For this a solvent system of (A) MeOH/H_2_O (85:15, v/v) and (B) MeOH/isopropanol (50:50, v/v) was used. Both eluents were modified by addition of small amounts of formic acid (0.12 %, v/v) and 14.8 M NH3aq (0.04%, v/v). The elution program was as follows: 95% A for 3 min, followed by a linear gradient to 40% A at 12 min and then to 0% A at 50 min. These latter conditions were maintained until 80 min. The settings for the electrospray ionization probe, operated in positive ion mode, were: capillary temperature, 300°C; sheath gas (N_2_) pressure, 40 arbitrary units (AU); auxiliary gas (N_2_) pressure, 10 AU; spray voltage, 4.5 kV; probe heater temperature, 50°C; S-lens 70 V. The Q Exactive mass spectrometer was calibrated within a mass accuracy range of 1 ppm using the Thermo Scientific Pierce LTQ Velos ESI Positive Ion Calibration Solution. The IPLs were analysed with a mass range of m/z 350–2000 (resolution 70,000 ppm at m/z 200). For data-dependent tandem MS/MS (resolution 17,500 ppm at m/z 200) fragmentation of the 10 most abundant ions was used (stepped normalized collision energy 15, 22.5, 30; isolation width, 1.0 *m/z*). Dynamic exclusion was used to temporarily exclude masses (for 6 s) to allow selection of less abundant ions for MS/MS.

The generated output data were analysed using Xcalibur Qual Browser Software (Thermo Fisher Scientific). Identification of IPLs was based on accurate mass measurements, retention time and MS^2^ fragmentation patterns, following established criteria ^65^. As different IPL classes exhibit variable ionization efficiencies, their measured peak areas (in response units) do not directly correspond to actual relative abundances. Nonetheless, this analytical approach allows for valid comparisons among samples processed and analysed in the same way.

### Protein expression and purification

Full length CdvB1 and CdvB2 were expressed and purified as in ^66^. Briefly, pOPINS expression vectors containing either 6His-SUMO-CdvB1 of *S. acidocaldarius* (residues 9-214 to capture the correct start codon) and 6His-SUMO-CdvB2 of *S. acidocaldarius* were transformed into *E. coli* C43(DE3) and grown in 2xTY medium with 30 µg/mL kanamycin. Expression was induced at OD600 ~0.6-0.8 using 0.5 mM isopropyl-β-D-thiogalactoside (IPTG) and grown for 4 h at 37°C before harvesting. All proteins were purified as follows. Steps up until dialysis were performed at room temperature, otherwise at 4°C. Cells were resuspended in buffer A (30 mM CHES; 150 mM NaCl; pH 9.5), with DNase, RNase, and protein inhibitor tablets (cOmplete EDTA-free, Roche). Cells were lysed by sonication and debris pelleted. Imidazole was added to filtered lysate up to 20 mM, and flowed over a 5 mL HisTrap (Cytiva). Protein was protein was eluted stepwise (50 mM, 100 mM, 200 mM, 300 mM, 500 mM, 1000 mM imidazole). 1 mM TCEP, ~1 mg homemade GST-tagged SENP1 ^67^ and 1 mL of pre-washed Glutathione Sepharose 4B resin (Cytiva) were added to the pooled fractions, and dialysed overnight against buffer A with 1 mM TCEP, at 4°C. The sample was flowed over a gravity column to remove GST-SENP1, and over a HisTrap collecting the flowthrough, removing uncleaved protein. All samples then went through a HiLoad 16/600 Superdex 75 pg column in buffer B (30 mM CHES; 400 mM NaCl; pH 9.5). Suitable fractions were pooled and concentrated.

### Liposome preparation

DOPC (18:1 1,2-dioleoyl-sn-glycero-3-phosphocholine, Avanti Research 850375), Sphingomyelin (egg, Avanti Research 860061), Cholesterol (ovine wool, Avanti Research 700000), DOPS (18:1 1,2-dioleoyl-sn-glycero-3-phospho-L-serine, Avanti Research 840035), PI (18:1 Phosphatidylinositol, Avanti Research 850149), PI(3P) (18:1 Phosphatidylinositol 3-phosphate, Avanti Research 850150), PI(4)P (18:1 Phosphatidylinositol 4-phosphate, Avanti Research 850151), PI(5)P (18:1 Phosphatidylinositol 5-phosphate, Avanti Research 850152), PI(3,4)P_2_ (18:1 Phosphatidylinositol 3,4-phosphate, Avanti Research 850153), PI(3,5)P_2_ (18:1 Phosphatidylinositol 3,5-phosphate, Avanti Research 850154), PI(4,5)P_2_ (18:1 Phosphatidylinositol 4,5-phosphate, Avanti Research 850155) were resuspended per manufacturer’s instructions and stored at −20° C. Lipid compositions described in the text and figure legend were mixed and dried under a stream of nitrogen. Residual solvent was removed by ROTOVAC for a minimum of 4 hours at 30° C. Lipid films were resuspended to a final concentration of 1mg/ml lipid in buffer C (50 mM MES(NaOH); 50 mM NaCl; pH 6.0) by vortexing. Unilamellar vesicles were prepared by freeze-thaw cycles (10x snap freezing and thawing at 37° C). Vesicles were stored at −20° C and used immediately upon thawing.

### Pelletting assay

The pelleting protocol was based on ^66^. Protein aliquots stored at −80C were quickly thawed and diluted to 0.6 mg/mL in buffer C. To pellet any pre-formed aggregates, the protein samples were centrifuged in a TLA-100 rotor (Beckman Coulter) for 30 min at 100,000 *g*, at 4°C. 6 µL of the supernatant was mixed with 30 µL of pre-formed liposomes and incubated at RT for 10 min. The samples were spun at 100,000 *g*, 4°C for 30 min. The pellets were briefly washed and resuspended in 36 µL of buffer C, and analysed together with supernatant samples on NuPAGE 4-12% Bis-Tris SDS-PAGE gels (Invitrogen).

### Statistics

All statistical analysis was performed in GraphPad Prism 10. Details of statistical tests are provided in the figure legends. Significance was defined as *P* ≤ 0.05. Significance levels: ^*^*P* ≤ 0.05, ^**^*P* ≤ 0.01, ^***^*P* ≤ 0.001 and ^****^*P* ≤ 0.0001.

## Acknowledgments

We gratefully acknowledge Anchelique Mets, Denise Dorhout, and Monique Verweij for their technical support in lipid analysis. We kindly thank Javier Espadas, Aurelien Roux, and Adolfo Saiardi for their discussions and comments on this study. We also thank the other members of the Moore Simons Consortium Group (Alex Bisson, John Mallon, Anja Spang, Josh Hamm, Sergei Ovchinnikov, Sirui Liu, Juan P. Karlusich) for their discussions and insight. We thank the Light Microscopy and Flow Cytometry facilities at the MRC-LMB, and all the core staff at the MRC-LMB. Finally, we thank all members of the Baum lab for helpful discussions and support.

A.C. was funded by an EMBO Postdoctoral Fellowship (ALTF_1041-2021) and a Marie Sklodowska-Curie Individual Fellowship (101068523). Y-W.K. was funded by an EMBO Postdoctoral Fellowship (ALTF 903-2021). B.B. received funding from the Medical Research Council - Laboratory of Molecular Biology (MC_UP_1201/27), Wellcome Trust (222460/Z/21/Z) and Paul G. Allen Frontiers Group (Allen Distinguished Investigator Award 2024). B.B. and L.V. received funding from the Life Sciences–Moore-Simons Foundation (735929LPI).

